# Relationships between competition sprint cycling power and endurance power are high across ALL durations and energetic pathways

**DOI:** 10.1101/2023.01.06.522987

**Authors:** Hamish Ferguson, Chris Harnish, Sebastian Klich, Kamil Michalik, Anna Katharina Dunst, Tony Zhou, J. Geoffrey Chase

**Author notes:** Corresponding author: Hamish Ferguson, Centre for Bioengineering, Department of Mechanical Engineering, University of Canterbury, Christchurch, New Zealand.

## Abstract

This study investigates associations between power at several durations to show inter- relationships of power across a range of durations in sprint track cyclists. The currently-accepted hypothesis peak power holds a near perfect relationship with sprint performance, and thus a near 1:1 slope with power at sprint durations up to 30-s, is tested. The equally well-accepted and complementary hypothesis there is no strong association with power over longer durations is also tested. 56 data sets from 27 cyclists (21 male, 6 female) provided maximal power for durations from 1-s to 20-min. Peak power values are compared to assess strength of correlation (R^2^), and any relationship (slope) across every level. R^2^ between 15-s – 30-s power and durations from 1-s to 20-min remained high (R^2^ ≥ 0.83). Despite current assumptions around 1-s power, our data shows this relationship is stronger around competition durations, and 1-s power also still shared strong relationships with longer durations out to 20-min. Slopes for relationships shorter durations were closer to a 1:1 relationship than longer durations, but closer to long- duration slopes than to a 1:1 line. The present analyses contradicts both well-accepted hypotheses, and the concept of peak power being a primary metric for sprint cycling, based on very strong relationships to power from durations commonly associated with oxidative energetic pathways, as well as short durations. This study shows the importance and potential of training durations from 1-s to 20-min over a preparation period to improve competition sprint cycling performance.

## Introduction

Sprint track cycling at Olympic Games and Elite World Championship levels is a subsection of cycle sport requiring a combination of peak speed, power, strength and short term endurance (speed endurance) [1]. While sprint cycling requires both a high level of strength and power, the nature of racing is actually more within the speed-endurance realm [2]. Unlike Olympic Weightlifting, powerlifting, or peak power field events, sprint cycling races range in durations from 15-s to 75-s depending on event [3-5]. Thus, competition demands on energy supply to the working muscles come from all three pathways: Adenosine Triphosphate Phosphocreatine (ATP-PCr), glycolytic, and oxidative [6-9]. Depending on time duration of maximal effort during sprint cycling events there are different contribution of these three energy systems [10].

Despite the physiological data showing reliance on both glycolytic and oxidative energetic pathways for durations associated with competition sprint cycling, the priority of training is focused on peak power [11], peak strength in the gymnasium [12], and maximal torque after the cycling-specific isometric resistance training [13]. The application of this approach has limited empirical evidence [3, 14]. There is thus a well-accepted hypothesis peak power is unrelated to power at longer, oxidative or endurance durations.

Stone et al. [14], show a relationship between various strength exercise measures and standing start performance (25-m – 333.33-m). However, this research lacks a comparison with longer durations to determine if the relationship stays strong as distance increases. Interestingly Dorel et al. [3], concluded peak power, relative to frontal area or pedalling frequency, was the strongest predictor of flying 200-m performance on a velodrome, however analysis of their full data, provided within the paper, using simple analysis shows a better relationship between measures of the actual flying 200-m performance, than with peak power measures. Moreover, peak power output related to body mass express the ability to accelerate [15].

Current models of sprint cycling performance appear inadequate to develop optimal performance in competition sprint cycling [1]. The reliance on peak power as a singular, simple, and main predictor of sprint cycling performance appears to be more about convenience than empirical evidence [15]. In particular, ergometer-based testing performed in the laboratory is widely used to assess this value, while successful performance in track-spring cycling is multi-factorial, incorporating physiological traits, tactics, and other specific skills [16].

The anaerobic speed reserve, developed to model track running performance for events from 10- s – 300-s, is based on maximal sprinting speed and maximal aerobic speed (the lowest running velocity at the *V*O_2MAX_ may occur) [17]. A cycling power meter is the most useful tool to determine relationships beyond peak speed/power and velocity/power at the *V*O_2MAX_, thus it is not surprising this model has been applied to road cycling sprinting, and aptly named the anaerobic power reserve (APR) [18]. However, the use of only two parameters to assess events up to 300s fails to account for the variety of energetic pathways, and accumulation of waste products, while competing in sprint cycling [19]. In regards to a suggestion by Leo et al. [20], the APR model may provide a useful concept to predict the power-duration relationship in the extreme exercise intensity domain, such as during sprint track cycling events. Current and future research studies should argue against the necessity to run multiple trials to predict performance given the ease of measuring actual performance [21], modelling the determinants from actual performance [22, 23], and to measure performance with various sensors, especially power output, during competition [24].

Investigations into the muscle fibre typology of cyclists from BMX, possessing the highest observed peak powers [25, 26], to professional road cyclists who have much lower peak powers in general [27-29] showed track sprint cyclists had a variable fast typology over lapping with both BMX and track endurance cyclists [30]. These data would suggest performance in sprint cycling is not tied solely on having muscle of the largest fast twitch fibre typology. However, applying the same analysis to swimmers saw no differentiation between muscle fibre typology in sprint swimmers, medium distance swimmers and long distance swimmers [31].

A question around modelling sprint cycling arises since the evaluation of actual performance with time, speed, power, and cadence data are easily accessible from performance. Moreover, the modelling becomes more feasible through the use of video data to assess the technical and tactical effectors of any given competition sprint race [21]. Due to the inconsistencies in the literature between measures of peak power and measures of peak sprint cycle racing performance, this research compares peak power in sprint cycling across a wider range of durations from 1-s to 20-min to assess the validity of assumptions in current training approaches.

Hence, this study investigates and quantifies associations between peak power and power at several durations from 1-s to 20-min to show inter-relationships of power across a range of training durations in sprint track cyclists. We hypothesise: (1) strong consistent correlation between sprint cycling power at 15-s and 30-s to power from 1-s out to 20-min; and (2) a slope closer to 1:1 will be seen for correlations between 15-s and 30-s power than for 1-s power. Both hypotheses contradict well-accepted belief and coaching practice in the field.

## Methods

### Participants

The online open-access, publicly available fitness depository www.strava.com [32], was used to access cycling data from training, testing and racing data from track cyclists in New Zealand. Track riders were identified competing and training at various track locations in New Zealand. Peak power metrics were harvested from riders who publicly posted their training and racing rides with power meter data publicly displayed. Sprint cyclists were identified by matching their Strava data against their performances at New Zealand national track cycling championships (www.cyclingnewzealand.nz [33]) and World Masters Track Cycling Championships (www.mastersla.com [34]). Strava and Cycling NZ age based championship data was used to ascertain weight, sex and age for the rider data. University of Canterbury Human Research Ethics Committee gave an exemption to seek ethics approval for the this research (2022/06/EX).

We harvested data from 27 junior, under 17 years old (U17): turn 15/16 years in year of competition, U19: turn 17/18, elite (turn 19-34), and masters (turn 35+ assignment in 5-year age groups) sprint track cyclists (Table 1). The classification of sprint track cyclists was made from the ability to place top 4 in the sprint in U17, U19, and Elite levels, top 4 in Keirin for U19, Elite and Masters, top 4 in time trial (500-m for women, 1000-m for men) in all grades, competing at Junior World level in sprint events, and top 4 placing in Masters World Championship events. Data from power meter measures were collected during testing, training, and racing sessions over a cross-section of male (21) and female (6) track cyclists.

**TABLE 1.** Participant data including this study, endurance (END) cyclists.

|  | **All:** Median [IQR] | **Male:** Median [IQR] | **Female:** Median [IQR] |
| --- | --- | --- | --- |
| **Data Sets** | 56 | 44 | 12 |
| **Participants** | 27 | 21 | 6 |
| **Age (years)** | 19 [16 - 39] | 25 [16 - 40] | 17 [16 - 19] |
| **Weight (kg)** | 76.5 [74.0 - 86.8] | 80.3 [75.0 - 88.0] | 64.0 [60.0 - 74.0] |
| **END Data Sets** | 144 | 128 | 16 |
| **END (Age)** | 26 [17 - 38] | 27 [17 - 39] | 19 [17 - 30] |
| **END (Weight)** | 70.0 [65.0 - 75.5] | 72.0 [67.0 - 76.0] | 57.0 [56.0 - 63.5] |

The final inclusion criteria required data for a range of durations spanning energy systems from ATP-phosphagen (1-6 s), oxygen-independent glycolysis (12-30s), and oxidative (over 40-s) energetic pathways [10]. Specifically: 1-s, 6-s, 12-s, 15-s, 18-s, 24-s, 30-s, 40-s, 50-s, 60-s, 90-s, 2-min, 3-min, 6-min, 12-min and 20-min, over a minimum of 3-months and maximum of 12- months of data. Data included maximal efforts over all durations from racing, testing and training sessions. To validate our data we used endurance cyclist data from previous research [2], to compare with our sprint cyclists data.

### Study Overview

Data collected from periods including testing, training and racing sessions, focused on sprint cycling competition performance was used. We measured 1-s, 6-s, 12-s, 15-s, 18-s, 24-s, 30-s, 40-s, 50-s, 60-s, 75-s, 90-s, and 2-min, 3-min, 6-min, 12-min, and 20-min maximal power for a training period of six to twelve months. First, this data was analysed to determine the relationships between all durations and two common durations in sprint cycling performance (15-s and 30-s). Second, the slope for the two sprint durations and all other durations was assessed and plotted compared to a 1:1 relationship (15-s:15-s and 30-s:30-s power).

The 15-s duration reflects the flying 200-m where the riders jump 100-m - 125-m from the 200- m mark on the track, the first lap of Team Sprint and the shortest duration match sprints take place over. Similarly, 30-s duration reflects a 500-m time trial, riding *2nd wheel* (2 rider in the starting line-up) in the team sprint, a long match sprint, and/or the Keirin event.

### Power Meter Data

All variables in this study are power meter data, measuring power in watts in 1-s intervals. An inclusion criteria for power meter data, was the use of a power meter model allowing regular calibration [35], and before each use a zero offset (reset torque measure to zero) was performed to account for changes in temperature. Collections of data were sent to the first author, and saved in the WKO5 (TrainingPeaks, Boulder, CO) athlete performance data analysis software. WKO5 was used to summarise peak power at selected durations.

### Analysis

The descriptive statistics including mean maximal power and interquartile range and statistical analysis was performed using Matlab version R2021a (The MathWorks, Natick, MA). Model quality was assessed by total least squares correlation coefficient, R^2^, to account for variability and error in both variables [36, 37]. A higher R^2^ value indicates a stronger relationship, and a better model and predictor. Model slope indicates the trade-off between power interval training and peak 1-s power. A Welch’s *t-*test for unequal sample sizes and variances was used to compare the differences in power between the sprint and endurance groups to ensure the study group were sprint cyclists in terms of power metrics [38].

## Results

### Assessing the Sprint Cohort

Table 2 shows the results of the Welch’s t-test between sprint and endurance cyclists from prior research [2], describing the difference between sprint and endurance groups for all bar the 30-s W/kg duration. Below the 30-s duration the sprint groups delivers higher power, and longer than 30-s, the endurance group deliver higher power for the 3-min and 20-min durations. These data indicate the group of national level sprint cyclist data collected were different from a group of national level endurance cyclists in a manner expected for sprint cyclists.

**TABLE 2:** Independent Samples T-Test between sprint and endurance for selected durations.

|  | Statistic | df | p |
| --- | --- | --- | --- |
| W/kg 1-s | 2.362 | 73.1 | 0.021 |
| W/kg 15-s | 2.163 | 70.0 | 0.034 |
| W/kg 30-s | 0.612 | 74.4 | 0.542 |
| W/kg 3-min | -6.591 | 64.4 | < .001 |
| W/kg 20-min | -10.447 | 68.5 | < .001 |

### Sprint Cycling Power Across Durations

Table 3 shows median and interquartile range peak power for all durations for all participants, and then delineated for male and female cyclists. It also shows the expected hyperbolic curve as power drops with an increase in duration, as well as expected differences between male and female athletes. Endurance cyclist power was included at selected intervals (boxed in table 3) to show the two groups are different, with endurance riders having lower power in short durations, and greater power at the longest durations, as expected.

**TABLE 3:** Mean maximal power (watts/kg) for all durations, median [interquartile range] for sprint cyclists. Boxed sections of the table showing 1-s, 15-s, 30-s, and 20-min power have values for both sprinters and in red for END or endurance cyclists to show the differences in power between these two groups.

|  | All Median [IQR] | Male Median [IQR] | Female Median [IQR] |
| --- | --- | --- | --- |
| **1-s PO** | 19.11 [16.61 – 20.39] | 19.43 [17.68 – 20.92] | 17.33 [16.08 – 17.95] |
| **END 1-s** | 17.71 [15.96 – 19.34] | 17.84 [16.39 – 19.48] | 15.61 [14.75 – 17.73] |
| **6-s PO** | 16.99 [14.95 – 18.90] | 17.71 [15.91 – 19.09] | 15.01 [14.57 – 17.03] |
| **12-s PO** | 15.41 [13.34 – 16.70] | 15.61 [14.09 – 16.89] | 13.83 [12.97 – 15.53] |
| **15-s PO** | 14.27 [12.91 – 15.93] | 14.63 [13.23 – 15.97] | 12.96 [12.12 – 14.27] |
| **END 15-s** | 13.22 [12.01 – 14.83] | 16.24 [14.91 – 18.08] | 12.07 [10.98 – 13.66] |
| **18-s PO** | 13.31 [11.97 – 15.08] | 13.53 [12.26 – 15.12] | 12.09 [11.27 – 13.28] |
| **24-s PO** | 11.97 [10.99 – 12.85] | 12.41 [11.10 – 13.40] | 11.10 [9.31 – 12.41] |
| **30-s PO** | 10.91 [9.91 – 12.00] | 11.02 [10.21 – 12.03] | 10.10 [8.40 – 11.40] |
| **END 30-s** | 10.61 [9.86 – 12.03] | 10.72 [9.98 – 12.14] | 9.97 [9.16 – 10.55] |
| **40-s PO** | 9.39 [8.23 – 10.31] | 9.49 [8.81 – 10.34] | 8.14 [7.67 – 9.34] |
| **50-s PO** | 8.47 [7.37 – 9.32] | 8.56 [7.77 – 9.50] | 7.56 [6.36 – 8.53] |
| **60-s PO** | 7.57 [6.61 – 8.48] | 7.68 [7.10 – 8.56] | 6.55 [5.99 – 8.04] |
| **75-s PO** | 6.64 [6.02 – 7.46] | 6.83 [6.12 – 7.53] | 6.27 [5.23 – 7.21] |
| **90-s PO** | 6.12 [5.56 – 7.05] | 6.22 [5.58 – 7.08] | 5.77 [5.14 – 6.93] |
| **2-min PO** | 5.53 [4.75 – 6.56] | 5.55 [4.87 – 6.57] | 5.16 [4.61 – 6.55] |
| **3-min PO** | 4.72 [4.35 – 5.83] | 4.73 [4.39 – 5.98] | 4.70 [4.32 – 5.47] |
| **END 3-min** | 6.14 [5.68 – 6.70] | 6.28 [5.79 – 6.76] | 5.65 [5.34 – 6.06] |
| **6-min PO** | 4.03 [3.60 – 4.92] | 4.04 [3.66 – 5.05] | 4.00 [3.58 – 4.66] |
| **12-min PO** | 3.69 [3.20 – 4.60] | 3.69 [3.21 – 4.68] | 3.60 [3.15 – 3.89] |
| **20-min PO** | 3.47 [3.01 – 4.12] | 3.53 [3.02 – 4.19] | 3.43 [2.90 – 3.62] |
| **END 20-min** | 4.76 [4.39 – 5.14] | 4.82 [4.43 – 5.24] | 4.45 [4.05 – 4.58] |

Table 4 presents the R^2^ values and slopes for 15-s and 30-s against power at all durations. The main result was the coefficient of determination (R^2^) was very high for all durations (R^2^ ≥ 0.87). In particular, correlations were very strong, or stronger, for durations longer than the 15-30-s durations most associated with sprint performance. Strong association was consistent (R^2^ ≥ 0.87) to the 20-min duration.

**TABLE 4.** R^2^ and slopes for 15 and 30 second power and power at all durations.

| **Time** | **R<sup>2</sup> 15-s** | **Slope 15-s** | **R<sup>2</sup> 30-s** | **Slope 30-s** |
| --- | --- | --- | --- | --- |
| **1-s** | 0.94 | 0.78 | 0.89 | 0.59 |
| **6-s** | 0.97 | 0.83 | 0.90 | 0.65 |
| **12-s** | 0.99 | 0.92 | 0.91 | 0.73 |
| **15-s** | **1.00** | **1.00** | 0.93 | 0.78 |
| **18-s** | 0.99 | 1.07 | 0.94 | 0.82 |
| **24-s** | 0.96 | 1.16 | 0.98 | 0.93 |
| **30-s** | 0.93 | 1.26 | **1.00** | **1.00** |
| **40-s** | 0.90 | 1.50 | 0.97 | 1.16 |
| **50-s** | 0.89 | 1.65 | 0.94 | 1.30 |
| **60-s** | 0.89 | 1.84 | 0.93 | 1.41 |
| **75-s** | 0.88 | 2.09 | 0.91 | 1.58 |
| **90-s** | 0.88 | 2.23 | 0.90 | 1.72 |
| **2-min** | 0.88 | 2.47 | 0.89 | 1.92 |
| **3-min** | 0.87 | 2.67 | 0.87 | 2.09 |
| **6-min** | 0.87 | 3.25 | 0.85 | 2.47 |
| **12-min** | 0.86 | 3.59 | 0.84 | 2.76 |
| **20-min** | 0.86 | 3.88 | 0.83 | 2.96 |

The slopes in Table 4 show all associations were increasingly further from the 1:1 line showing a reducing, but still strong, trade-off as duration increases, as expected. More specifically, a 1.1x or 10% change in peak power at longer durations will yield a smaller change in 1-s power than a similar 10% change in power for shorter durations, as the slopes in Table 4 decrease as duration. Slopes greater than 1.0 in Table 4 indicate the watts in 1-s gained per 1.0 Watt gain in power at the specified duration.

Similarly, Figure 1 shows the matrix of correlation coefficients comparing all power durations. Each power was perfectly correlated to itself. The minimum values are all R^2^ ≥ 0.82, showing there are no distinguishable or large differences in correlations between power across all durations. Table 4 and Figure 1 fully confirm our first hypothesis.

**Figure.**
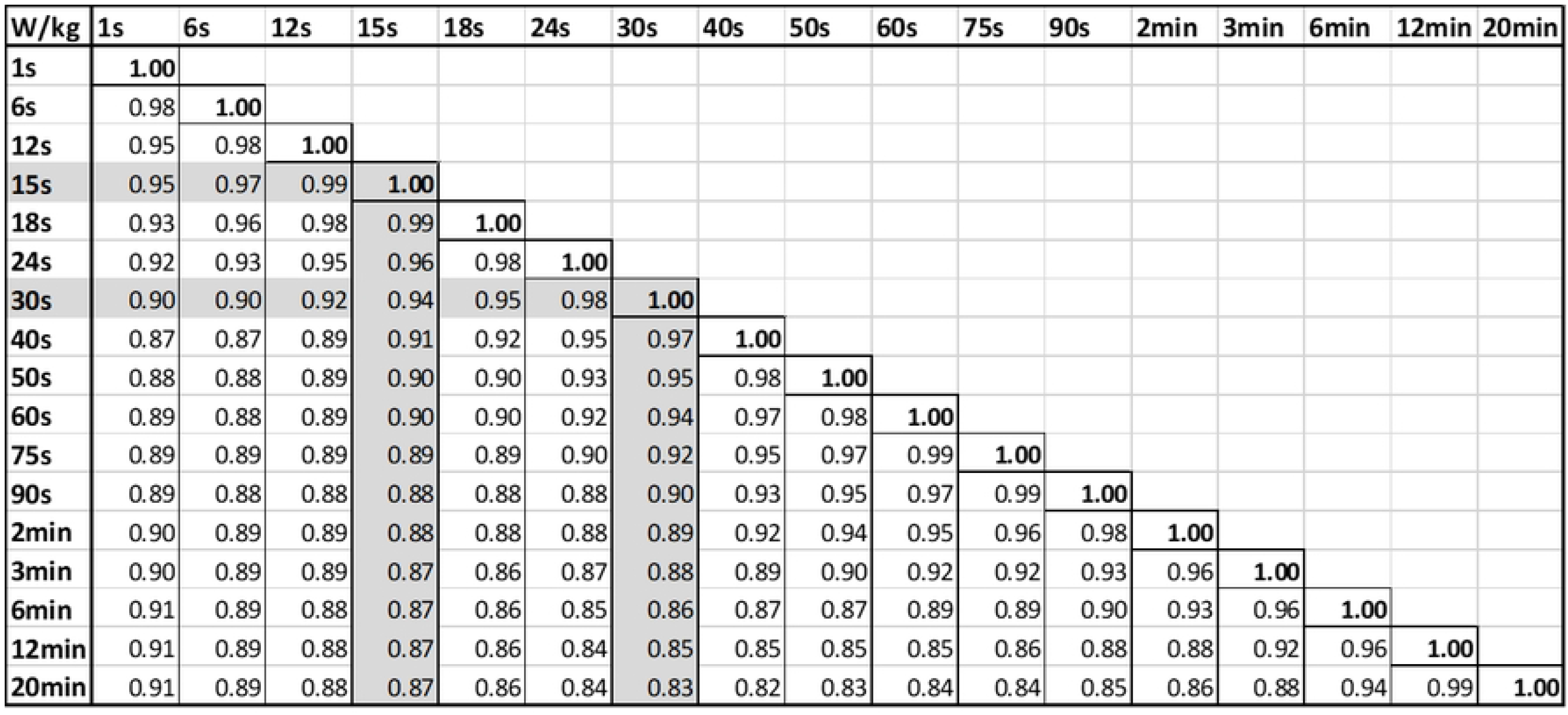

## Discussion

The purpose of this study was to compare peak power in sprint cycle racing for a wide range of durations. We hypothesised there would be strong correlation across all ranges of power measured from 1-s to 20-min, contrary to current coaching belief and practice. In particular, we tested these relationships, particularly to two common sprint cycling durations (15-s and 30-s) and durations longer durations where oxidative energy supply predominates.

This research is the first to demonstrate relationships between competition sprint power with power durations up to 20-min, and suggests oxidative energetic pathways are strongly associated with power output and measures of performance in competition sprint cycling. This study provides data and guidelines for sprint coaches on how to use this research to guide training through different periods leading into key competitions.

The data for a group of national level sprint cyclists was different to a group of national level endurance (road and track endurance) cyclists (Table 2). Data comparing power indicate the strongest relationships are found with the durations around the 15-s and 30-s power durations reflecting performance in actual sprint competition (Figure 1). Peak 1-s power is strongly related to sprint competition power, but this relationship stays strong all the way out to 20-min power (Figure 1), showing training these durations is not detrimental to competition sprint cycling performance.

Slopes for all power durations reiterate the strongest relationships are around 15-s and 30-s power (Figure 1). The 1-s power slope has the same slope as 2-min power, which shows all the durations between these values are important to train to influence sprint performance. This outcome is also contrary to current coaching practice and belief in the field, and these results provide significant evidence supporting a different approach.

In particular, while 1-s is a very strong predictor of performance for both the 15-s and 30-s durations common to actual sprint competition, 6-s, 12-s, 18-s, and 24-s are stronger predictors for 15-s power. Similarly, 6-s, 12-s, 15-s, 18-s, 24-s, 40-s, 50-s, 60-s, 75-s and 90-s power are all better predictors of 30-s power performance than 1-s power. It also indicates longer time intervals should be used in power-duration modelling, not only the 1-s peak power during the ∼10-s – 15-s sprint effort, as also suggested by Sanders and Heijboer [18].

Our findings indicate the slopes for all durations do not achieve the 1:1 relationship, with all tested durations reflecting actual sprint cycling competition durations showing slopes closer to 1:1 than 1-s power. For example, looking at 15-s power, 6-s, 12-s, 18-s and 24-s power all have slopes closer to 1:1 than 1-s power. For 30-s power, 6-s, 12-s, 15-s, 18-s, 24-s, 40-s and 50-s power have a closer slope to 1:1 than 1-s power. These data support Hypothesis 2. For practical application, each position from Team Sprint should be tested in different specific ways: wheel 1 15-20-s, wheel 2 30-35-s and wheel 3 using test lasting 40-s. This approach is based on different metabolic demands and energy systems contribution [39].

Finally, our data indicate a strong relationship between sprint cycling power durations of 15-s and 30-s, and durations primarily fuelled by oxidative energetic pathways out to 20 minutes. Strong R^2^ values extend all the way out to 20-min durations in support of this interpretation. This outcome shows a sprint cyclist can deliver a high 20-min power, relative to other sprint cyclists, and not relative to endurance cyclists who display higher 20-min power outputs and lower 1-s, 15-s & 30-s power (Table 4). While data from road cyclists showing power at the end of a long road race was lower than their peak power, it confirms prior exercise can influence peak power delivery [29]. Improving aerobic energetic pathways is also important for sprint cyclists to enhance recover between maximal efforts in training sessions and competitions [40].

Taken together these data provide a more complete picture of competition duration sprint cycling power, and their relationships with power durations fuelled not only by the ATP-PCr system, but also the glycolytic and oxidative energetic pathways. The data also reflect the neuromuscular demands of competition duration sprint cycling performance, where the best relationships are seen between competition similar durations rather than 1-s power. Finally, the data may also reflect the nature of sprint cycling competition where the ability to perform repeatedly and recovery from high intensity intermittent exercise are also a necessity.

This overall outcome challenges the notion of an anaerobic speed or power reserve and supports the development of performance to meet the actual demands of sprint cycling performance over common durations seen in racing. This type of analysis can be used to profile cyclists in terms of developing the capacity to race maximally for 15-s – 30-s, such as in the sprint and Keirin events, where the exact duration of competition remains unknown depending on the group dynamics of each race [1, 41, 42]. The need to raise capacity would be seen in those riders who sit above the line of best fit, possessing high power output, and who would potentially benefit more from increasing the duration they can sustain their power and enhanced repeatability of performance with all championship sprint events requiring multiple efforts to medal.

In particular, how far away from the line a powerful rider is located at a given point in a training period could indicate the best stimulus to improve their capacity to hold good power for a long duration. The data showing a 20-min effort still has a strong relationship with 15-s and 30-s sprint performance suggests these stimuli could vary, particularly in different training phases in preparing for key events or a season. Equally, a rider below the line of best fit would potentially benefit from increasing their power targeting on shorter more intense efforts, and again the further from the line the shorter the duration in training to try and increase their power relative to actual sprint cycling performance durations.

Limitations to this data are the group is based on National level sprint cyclists. While our data is different to the data of endurance cyclists (Tables 4 and 5), it is still likely our participants will conduct more endurance riding, and do track endurance racing and road cycling events than a high-performance group. The question is whether this difference reflects tradition or a thorough analysis of the actual demands of competition sprint cycling performance. Due to challenges with Covid-19 we elected to harvest data from a publicly available open source. We feel the use of data from this natural setting potentially enhances the quality of the research as riders are performing for the sake of performance with no other external influence, rather than just for research testing, which could be seen as a chore.

We also need to comment on the use of power and highlight how velocity over the whole sprint wins sprint cycling events. With proper use of the track, pacing over the sprint, position on the bicycle, position on the track: where riding the black measurement line is the shortest distance on the track (bankings allow a rider to use gravity to build speed rather than delivering power), positioning in the race (1 or 2 in a sprint, or 1 – 6 in a Keirin) and how a rider passes uses sound tactics. All offer means of increasing or maintaining velocity which may not be reflected in power meter data. Velocity can be maintained while power is dropping during a sprint [43].

This data is based of sprint cycling training and racing over a 3-6 month period of national level sprint cyclists, where it is likely 20-min peaks were performed early in a general preparation phase [44]. Shorter duration efforts are more likely to be performed in the lead up to competition. This commonly reflects the periodisation of training focusing on three distinct periods of general preparation, specific preparation and peaking for key events [44]. Some training programmes utilise a reverse periodisation approach referred to as short to long, focusing on peak power [45]. However, the data presented here suggests a focus on 1-s is potentially to the detriment of actual performance when developing power around competition times has a closer relationship.

Future research should focus on development of power profiles specific to sprint cyclists to determine whether they should focus on building capacity to ride hard for common durations of sprint cycling performance or whether they should increase their power over these durations. Elite level sprint cycling programmes should experiment on focusing on both capacity, as well as current practices based around building 1-s power. With strong relationships between sprint cycling power and 20-min power experimenting with longer efforts would seem a potential area to develop a more complete sprint cyclist.

## Conclusions

In a study comparing maximal efforts for durations supplied by all three energetic pathways and reflecting different neuromuscular demands we saw strong relationships between durations from 1-s - 20-min power in national level sprint cycling competitors. The strongest relationships were seen with durations around two key durations in sprint cycling performance: 15-s and 30-s power. Performance at durations of 2-min – 20-mins, primarily fuelled by oxidative pathways, were strongly correlated and did not diminish performance of either 15-s or 30-s. We also contend the development of a sprint cyclist should focus on first, the development of power from 1-s - 20-min in a general preparation phase, and then to more specific power around the 15-s – 30-s duration and peaking to focus on the demands of the events at a key event.

## Author contribution

HF conceived the study, performed the data collection and analysis, and wrote the original draft. TZ wrote the code used to analyse the data. JGC and CH performed the original edits to the manuscript. AKD, SK and KM all performed edits to the manuscript and all contributed to all aspects of the paper.

## Ethics statement

Open source data was used, so ethics was not required for this study. This exemption was approved by the University of Canterbury, Christchurch, New Zealand (2022/06/EX).

## Data Availability Statement

Data is available at: https://osf.io/2y3kr/

